# The *TERT* promoter is polycomb-repressed in neuroblastoma cells with long telomeres

**DOI:** 10.1101/2022.05.18.492493

**Authors:** Mindy Kim Graham, Beisi Xu, Christine Davis, Alan K. Meeker, Christopher M. Heaphy, Srinivasan Yegnasubramaian, Michael A. Dyer, Maged Zeineldin

**Affiliations:** Department of Radiation Oncology and Molecular Radiation Sciences, Johns Hopkins University School of Medicine, Baltimore, MD, 21205; Center for Applied Bioinformatics, St. Jude Children’s Research Hospital, Memphis, Tennessee 38105, USA; Department of Pathology, Johns Hopkins University School of Medicine, Baltimore, MD, 21287; Department of Oncology, Johns Hopkins University School of Medicine, Baltimore, MD, 21287; Department of Urology, Johns Hopkins University School of Medicine, Baltimore, MD, 21287; Department of Medicine, Boston University School of Medicine, Boston, MA, 02215; Department of Developmental Neurobiology, St. Jude Children’s Research Hospital, Memphis, Tennessee 38105, USA; Howard Hughes Medical Institute, Chevy Chase, Maryland 20815, USA; Department of Ophthalmology, University of Tennessee Health Science Center, Memphis, Tennessee 38163, USA

## Abstract

Acquiring a telomere maintenance mechanism is one of the hallmarks of high-risk neuroblastoma and commonly occurs by expressing telomerase (*TERT*). Telomerase-negative neuroblastoma has, characteristically, long telomeres and most utilize the telomerase-independent alternative lengthening of telomeres (ALT) mechanism. Conversely, no discernable telomere maintenance mechanism is detected in a fraction of neuroblastoma with long telomeres, representing a phenomenon referred to as ever-shorter telomeres. Here, we show that, unlike most cancers, DNA of the *TERT* promoter is broadly hypomethylated in neuroblastoma. In telomerase-positive neuroblastoma cells, the hypomethylated DNA promoter is approximately 1.5-kb in length and is bound by hypermethylated *TERT* gene body and upstream intergenic sequences. The *TERT* locus shows active chromatin marks including H3K4me3, H3K27Ac, H3K14Ac, RNA PolII and BRD4 with low enrichment for the repressive mark, H3K27me3. Strikingly, in neuroblastoma with long telomeres, the hypomethylated region spans the entire *TERT* locus, including multiple nearby genes with enrichment for the repressive H3K27me3 chromatin mark. Furthermore, subtelomeric regions showed enrichment of repressive chromatin marks in neuroblastomas with long telomeres relative to those with short telomeres. These repressive marks were even more evident at the genic loci, suggesting a telomere position effect. Inhibiting H3K27 methylation by the EZH2 inhibitor, Tazemetostat, induced the expression of *TERT*, particularly in cell lines with long telomeres and H3K27me3 marks in the promoter region. Taken together, these data suggest that epigenetic regulation of *TERT* expression differs in neuroblastoma depending on the telomere maintenance status, and H3K27 methylation is important in repressing *TERT* expression in neuroblastoma with long telomeres.

## Introduction

Telomeres are tandem hexanucleotide repeats located at eukaryotic chromosomal termini that in most normal somatic cells shorten with every replication cycle^1^. Stem and progenitor cells express a reverse transcriptase called telomerase (TERT) that utilizes a long non-coding RNA template (*TERC*) to add telomeric repeats specifically to chromosomal ends to maintain telomeres^1^ and prevent telomere-induced senescence and apoptosis^2^. Expression of *TERT* is strongly repressed in most somatic cells, limiting the number of replications that a cell can go through (Hayflick limit) and protecting against malignant transformation^3^. To overcome this key replicative barrier, the majority of cancers express *TERT* to maintain telomere lengths^4^. However, approximately 4-11% of cancers rely on a telomerase-independent recombination-mediated pathway termed Alternative lengthening of telomeres (ALT) to maintain telomere lengths^5^. In some rare cases, cancer cells with long telomeres are negative for both telomerase and ALT, suggesting either an undefined telomere maintenance mechanism, or lack of a mechanism^6,7^.

Neuroblastoma is a pediatric tumor that arises from the developing sympathetic neurons and is one of the leading causes of cancer-related death in children^8^. The acquisition of a telomere maintenance phenotype defines high-risk neuroblastoma and failure to maintain telomere lengths may lead to spontaneous regression of the tumor^9^. Telomere maintenance in neuroblastoma is controlled by genetic and epigenetic factors through three mutually exclusive mechanisms^10,11^. First, amplification of the oncogene *MYCN*, which is the most frequent genetic alteration seen in high-risk neuroblastoma^12–14^. MYCN is a transcriptional factor that induces transcriptional and epigenetic reprograming^15,16^ and *TERT* is a known MYCN target^17^. Second, in a subset of high-risk neuroblastoma cases, rearrangement of the *TERT* locus on chromosome 5 results in a juxtaposition of a proximal enhancer and *TERT* that consequently induces a high level of *TERT* expression, a process termed “enhancer hijacking”^11^. Finally, *ATRX* mutations, which are strongly associated with ALT, are frequently detected in high-risk neuroblastoma, especially in adolescents and young adults^16,18^. ATRX is a chromatin remodeler, and its mutations are associated with wide-spread epigenetic alterations^16,19-21^. Taken together, these observations suggest that epigenetic factors play an important role in telomere maintenance and the modulation of *TERT* expression in neuroblastoma.

DNA methylation and post-transcriptional histone modifications are two major epigenetic mechanisms that control gene expression^22^. Although extensively studied, the significance of the methylation of CpG islands in the *TERT* locus is unclear^23^. Proximal promoter methylation is generally considered a repressive mark that limits the accessibility of genes to transcriptional factors^22^. Paradoxically, *TERT* promoters are hypermethylated in telomerase-positive cancers and hypomethylated in most untransformed cells, suggesting that methylation is associated with telomerase expression in cancer^24,25^. However, many studies have consistently shown that indeed promoter DNA methylation inhibits *TERT* expression^26,27^.

In addition to DNA methylation, other epigenetic changes including post-transcriptional histone modifications control gene expression, with different combinations of chromatin marks defining distinct functional activities in the genome^28^. Acetylation of histone H3 lysine at positions 9 and 27 (H3K9Ac, H3K27Ac) and trimethylation at amino acid 4 (H3K4me3) reduce chromatin packing and permit the binding of transcription factors, thereby marking actively expressed genes. Conversely, H3K9me3 and H3K27me3 are associated with closed heterochromatin and considered repressive epigenetic marks^22^. These chromatin marks are installed and maintained by large multi-subunit protein complexes. For example, the polycomb repressive complex-2 (PRC2) plays an essential role in methylating H3K27 to ensure silencing of target genes^22,29^.

Recently, we characterized the epigenetic landscapes in a wide range of neuroblastoma cell lines, orthotopic patient-derived xenografts, and cancer tissues taken at time of autopsy^16^. In these samples, we performed whole genome bisulfite sequencing and ChIP-Seq for eight different chromatin marks and the epigenetic regulators: BRD4, RNA polymerase PolII, and the transcription regulator CTCF. These data were compiled into 18 different epigenetic states using Hidden Markov Modeling (HMM). Additionally, we mapped the genomic distribution of MYCN binding sites in the *MYCN*-amplified cells^16^. Here, we used these data to comprehensively examine the epigenetic regulation of the *TERT* promoter region in neuroblastomas. We found that, similar to normal tissues of developing adrenal medulla, the *TERT* promoter is hypomethylated in all neuroblastoma cells we studied, independent of telomere maintenance status. However, in neuroblastoma cells with long telomeres, including ALT-positive cells, we observed that the hypomethylation of the *TERT* promoter extended throughout the *TERT* promoter CpG island. In contrast, the *TERT* gene body is hypermethylated in telomerase-positive neuroblastoma cells. We also found that telomerase-positive neuroblastoma cells have the active chromatin marks H3K9Ac and H3K27Ac in the *TERT* promoter, while the *TERT* promoter of neuroblastoma cells with long telomeres are silenced by H3K27me3, suggesting polycomb repression of the *TERT* locus. Indeed, pharmacological inhibition of EZH2, the enzymatically active subunit of PRC2, resulted in an increase in *TERT* expression, suggesting de-repression of the *TERT* locus. Altogether, these data contextualize the epigenetic regulation of the *TERT* locus in high-risk neuroblastoma based on differing telomere maintenance mechanisms.

## Results

### *TERT* promoter CpGs are hypomethylated in neuroblastoma

To explore the epigenetic control of *TERT* expression, we leveraged our previously generated whole genome bisulfite sequencing (WGBS) analysis of neuroblastoma^16^. In our prior work, we comprehensively characterized the epigenetic marks of eight different cancer autopsies, seven orthotopic patient-derive xenografts (O-PDX) derived from these cancer autopsies, as well as eight additional neuroblastoma cell lines. Together, these 23 neuroblastoma samples represent the full spectrum of genetic alterations observed in high-risk neuroblastoma. Here, we analyzed DNA methylation at the *TERT* promoter and found approximately 1.5-kb of DNA hypomethylated region spanning from nucleotides −694 to +758 relative to the *TERT* gene transcription start site and overlapping the *TERT* promotor region in all neuroblastoma samples (Figure 1). This hypomethylated region surrounding the *TERT* transcriptional start site is also observed in the fetal adrenal medulla (Figure 1). This is particularly relevant as the fetal adrenal medullae contain progenitors of sympathetic ganglion cells and neuroblastoma is thought to originate from these progenitor cells that have undergone differentiation arrest.

**Figure 1:**
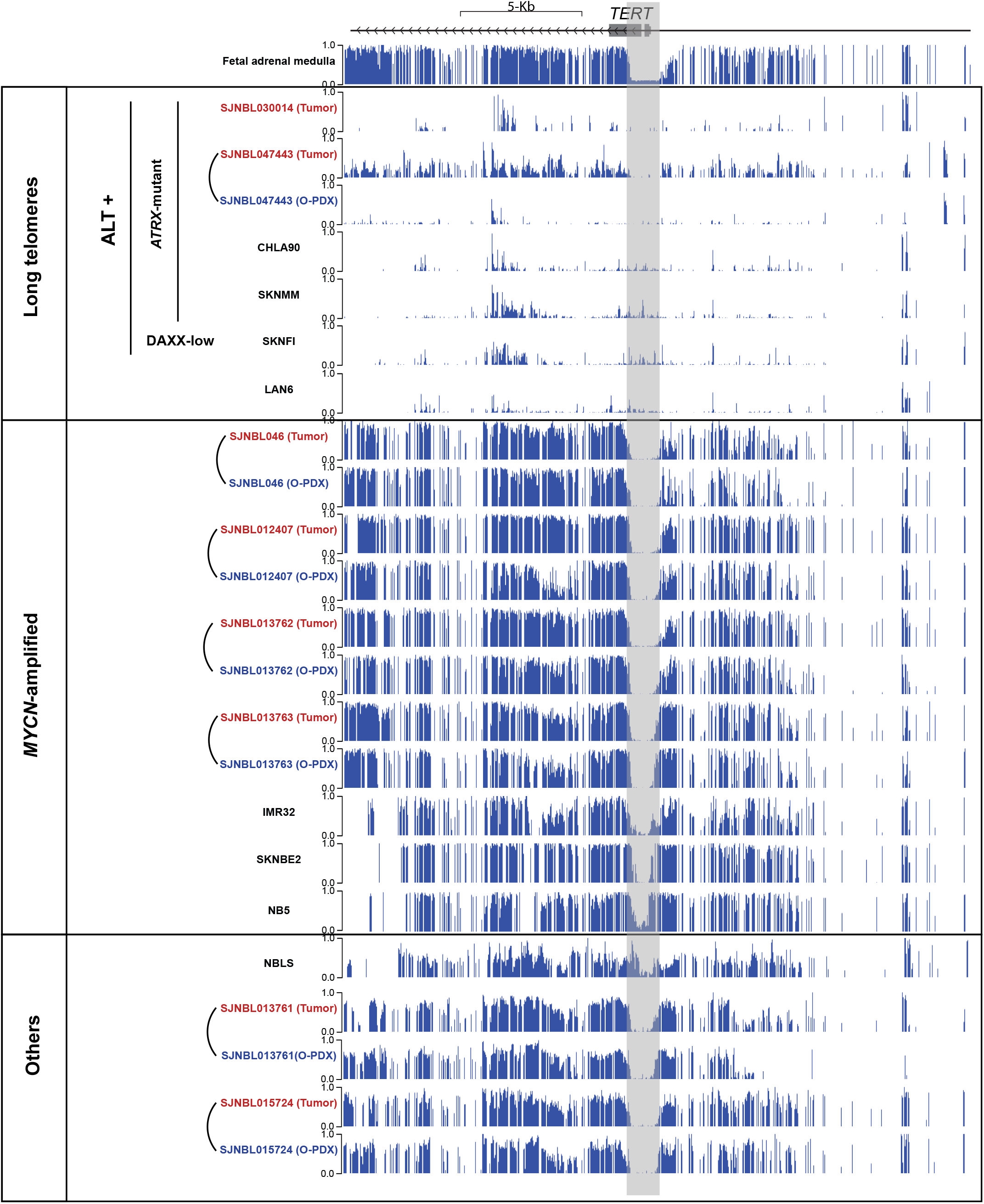
DNA hypomethylation of the TERT locus in neuroblastoma cell lines, tumors and O-PDX. **A)** Alignments of DNA methylation from whole genome bisulfite sequencing for the *TERT* locus in neuroblastoma cell lines (black), autopsy (red) and O-PDX (blue). Tumor autopsies and O-PDX that are derived from the same patients are connected by a curved line. The hypomethylated region shaded in gray starts −694 bases from the TERT transcriptional start site (TSS) and ends 758 bases after the TSS.

It was previously shown that the *TERT* proximal promoter is comprised of a core promoter located around the transcription start site (TSS, from +51 to −217), and a hypermethylated 433-bp region immediately upstream to the core promoter referred to as the *TERT* hypermethylated oncological region (THOR)^17,30^. As shown in Figure 2, the DNA hypomethylated region surrounding the TSS of *TERT* in neuroblastoma cells overlaps with this previously described core promoter. Moreover, in *MYCN*-amplified neuroblastoma, we observed MYCN occupation in this same hypomethylated region. Together, these data suggest that the *TERT* promoter hypomethylated region in neuroblastoma plays a role in regulating *TERT* expression.

**Figure 2:**
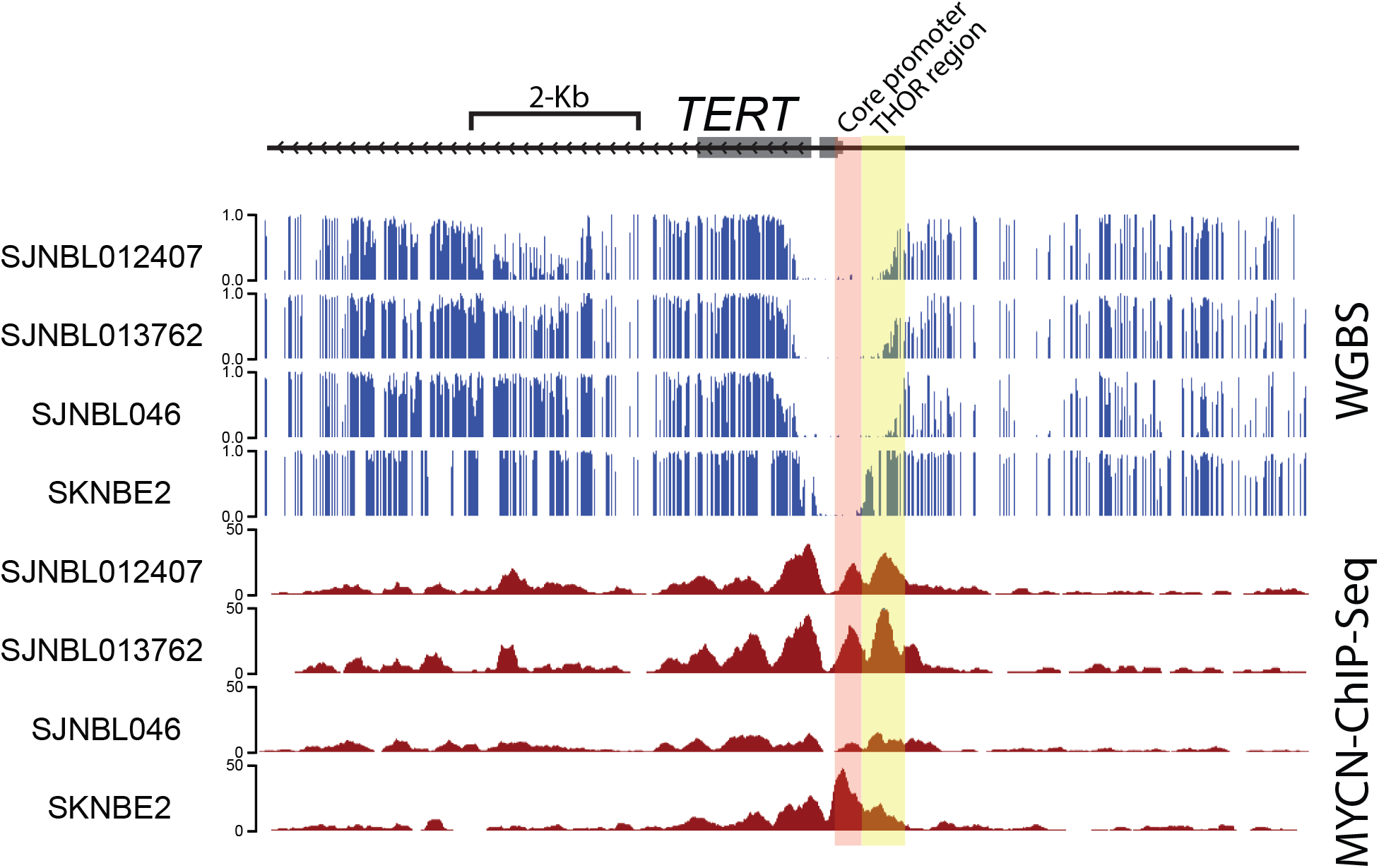
MYCN-binding sites overlap with the hypomethylated region in the *TERT* promoter. Alignments of DNA methylation from whole genome bisulfite sequencing (WGBS) and MYCN-ChIP-Seq in three MYCN-amplified neuroblastoma O-PDX models and cell line (SKNBE2) showing that MYCN binding overlaps with the hypomethylated region in the *TERT* promoter. The core promoter is shaded in red and THOR region is shaded in yellow.

### *TERT* promoter hypomethylation is expanded in neuroblastomas with long telomeres

The neuroblastoma samples were previously evaluated for *ATRX* and *DAXX* mutations, both of which are strongly associated with ALT, as well as *MYCN* amplification, which is observed in telomerase-dependent/ALT-negative neuroblastomas^16^. We expanded the characterization of telomere maintenance mechanism status of the eight neuroblastoma cell lines by performing telomere-specific FISH, telomere qPCR, and the C-circle assay, and measuring the relative expression of the telomerase catalytic subunit *TERT* and the RNA template *TERC* (Figure 3). Telomere-specific FISH staining revealed ultrabright telomeric foci, consistent with long telomere lengths, in SKNMM, CHLA90, SKNFI and LAN6, but not in NB5, SKNBE2, IMR32, and NBLS (Figure 3A). The relative quantity of telomere repeats assessed by real-time qPCR (Figure 3B) corresponded with the cell lines identified as having relatively longer telomeres. Partially single-stranded extrachromosomal circular DNA containing C-rich telomeric-repeat sequences, called C-circles, are enriched in ALT-positive cancer^31^. To measure C-circle levels in our neuroblastoma cell lines, we used a rolling circle amplification-based assay^32^. Neuroblastoma cell lines with short telomeres were C-circle negative, while cell lines with long telomeres were positive for C-circles, except for LAN6 (Figure 3C). *TERT* expression was consistently elevated in neuroblastoma cell lines with short telomeres, while neuroblastoma cell lines with long telomeres had extremely low levels of *TERT*. Taken together, these results confirm that SKNMM, CHLA90 and SKNFI are indeed ALT-positive, as they possess the hallmarks characteristic of ALT-positive cancers: long telomeres, very little to no expression of telomerase, and the presence of C-circles. Notably, SKNMM and CHLA-90 carry *ATRX* mutations^16,33,34^, which are tightly associated with ALT and detected in a significant fraction of high-risk neuroblastoma found in adolescents and young adult patients^16^. Although SKNFI has intact ATRX, the expression of DAXX, an established ATRX binding partner and ALT suppressor, is low^16,35^. ALT-associated *DAXX* mutations are observed in pancreatic neuroendocrine tumors but rarely in neuroblastoma^36^. Although LAN6 shares many features observed in ALT-positive cancers, the uniformity of telomere lengths, the absence of C-circles, and intact *ATRX* and *DAXX*^16^ suggests that it is neither ALT-positive nor telomerase-dependent, and may be similar to cancers previously described in the literature as ever-shorter telomeres^6^. All neuroblastoma cell lines in this study expressed *TERC*, with long telomere cell lines tending towards lower expression compared to short telomere cell lines (Figure 3E). We also confirmed *MYCN* amplification status in SKNBE2, IMR32 and NB5 cells using FISH (Supplementary Figure S1).

**Figure 3:**
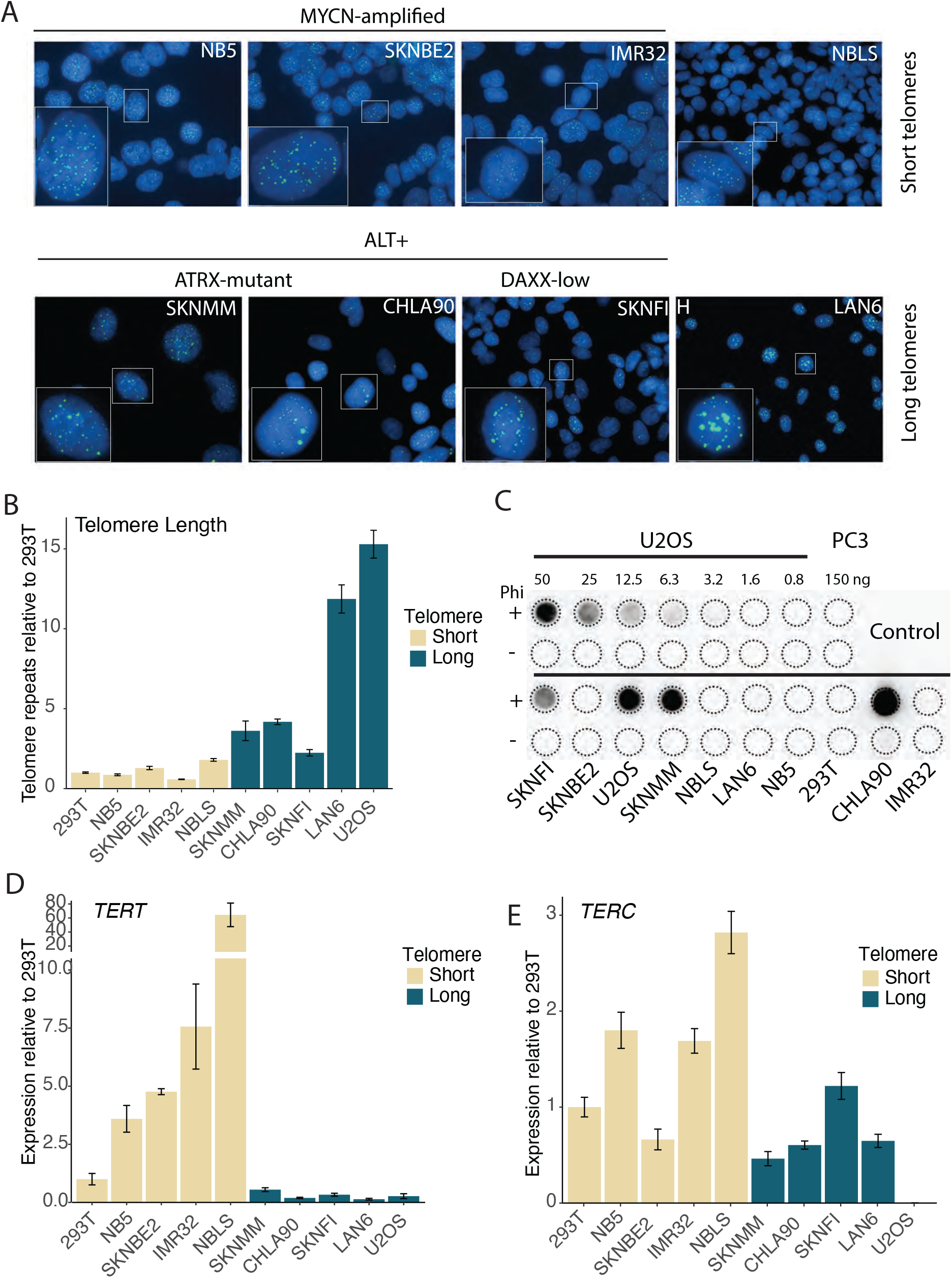
Molecular characterization of telomere maintenance in neuroblastoma cell lines. Telomere maintenance mechanism was assessed in eight neuroblastoma cell lines used in this study: MYCN-amplified (NB5, SKNBE2, IMR32), moderate level of MYCN without amplification (NBL-S), ATRX-mutant (SKNMM, CHLA90), low-DAXX (SKNFI) and ever-shorter telomere (LAN6) neuroblastoma cells. **A)** Representative images of Telomere FISH (green) showing brighter signal in cell lines with long telomeres. **B)** Telomeric repeats in neuroblastoma cell line relative to those in the telomerase positive 293T cells using qPCR. Telomerase negative, ALT-positive U2OS cells were used as a positive control for long telomeres. **C)** Representative blot of C-circle assay. U2OS cells were used as a positive control and PC3 cells were used as a negative control. The amount of DNA in nanograms (ng) are indicated on the blot. All reactions were done in the presence or absence of the Phi 29 (Phi) DNA polymerase. RT-qPCR was performed in neuroblastoma cell lines to measure **D**) *TERT* and **E**) *TERC* expression relative to the telomerase-positive 293T cell line. Gene expression was normalized to *HPRT* expression. The U2OS cell line was used as an ALT-positive control.

When we stratified the neuroblastoma cell lines by short versus long telomeres, a striking pattern emerged when we evaluated the methylation status of the *TERT* promoter CpG island. In neuroblastomas with long telomeres, the hypomethylated region extended beyond the proximal promoter region to include the entire *TERT* promoter CpG island, and multiple nearby genes (Supplementary Figure S2). Moreover, the O-PDX models and patient autopsies with *ATRX* mutations confirmed this pattern of extended DNA hypomethylation beyond the *TERT* promoter CpG island (Supplementary Figure S2). Taken together, these data suggest that telomerase-independent cancer cells with long telomeres are associated with repressed *TERT* expression and hypomethylation of the *TERT* locus in neuroblastoma.

### Neuroblastoma cells with long telomeres are enriched for repressive histone marks at the *TERT* locus

Histone tail modifications play key roles in regulating gene expression^22^. We examined the chromatin marks at the *TERT* locus and found that neuroblastoma cells with *MYCN*-amplification and short telomeres are enriched for chromatin marks indicative of open chromatin including H3K4me3, H3K27Ac, H3K14Ac, RNA PolII and BRD4 at the *TERT* promoter (Figure 4,A, B). These activating epigenetic marks overlap the region of DNA hypomethylation in the proximal *TERT* promoter, supporting a functional correlation between both the histone and DNA epigenetic marks. We also detected a low level of enrichment for the repressive mark H3K27me3 in the majority of the *MYCN*-amplified neuroblastoma samples, including the O-PDX sample (SJNBL046) which displayed a relatively high level of H3K27me3 (Supplementary Figure S3). Previously, we employed Hidden Markov modelling to bin epigenetic marks into 18 different epigenetic states (ChromHMM)^16^. The co-existence of repressive and active marks at the promoter region corresponds to our ChromHMM state 7 (bivalent promoter)^16^, which is frequently observed in some genes poised for rapid activation, especially during development^37^. Indeed, the proximal *TERT* promoter in fetal adrenal medullary cells showed this bivalent chromatin mark, with the enrichment of both H3K4me3 and H3K27me3 (Supplementary Figure S3). However, because we are unable to ascertain allele-specific epigenetic marks, this bivalent state may also represent monoallelic control of *TERT* expression, with one active allele and another transcriptionally silent allele.

**Figure 4:**
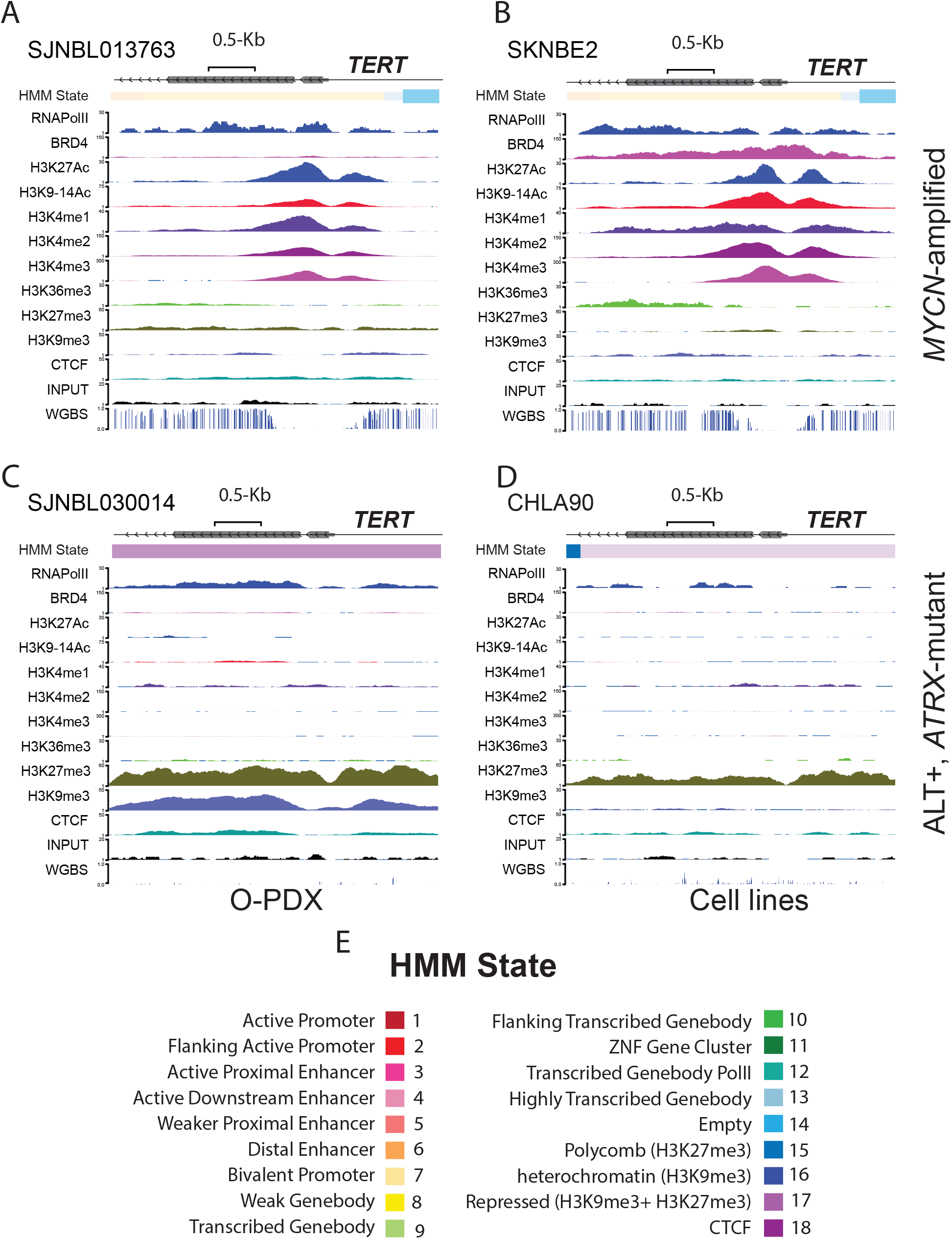
Active chromatin marks in MYCN-amplified neuroblastoma and repressive chromatin marks in neuroblastoma cells with long telomeres. **A-B** ChromHMM states,all ChIP-Seq tracks and whole genome bisulfite sequencing (WGBS) for MYCN-amplified O-PDX (SJNBL013763, **A**) and cell line (SKNBE2, **B**) showing active marks in the *TERT* locus. **C, D** ChromHMM states,all ChIP-Seq tracks and whole genome bisulfite sequencing (WGBS) for *ATRX*-mutant, ALT positive O-PDX (SJNBL030014, C) and cell line (SKNMM, D) showing repressive marks in the *TERT* locus. The color codes for different ChromHMM states are shown in **E.**

In contrast to the *MYCN*-amplified neuroblastomas, the ALT-positive *ATRX* mutant neuroblastoma autopsies and O-PDX tumors, as well as the cell lines with long telomeres, showed repressive chromatin marks, especially H3K27me3 in the same region of the proximal *TERT* promoter (Figure 4C, D). This H3K27me3 repressive mark overlaps the extended DNA hypomethylated region observed in neuroblastomas with long telomeres including ALT-positive tumors. The *TERT* promoter tended towards ChromHMM 17 state (Figure 4), which is characteristic of repressed regions of the genome. Consistent with the active and repressed epigenetic states segregated by short versus long telomeres, *TERT* expression was elevated in cells lines with active chromatin marks and short telomeres compared to cell lines with long telomeres and repressive marks (Figure 3D).

Long telomeres may loop back over chromosomal ends, interacting with many genes over long distances and repressing their expression^38–40^. Since *TERT* is close to the chromosome 5p terminus, it suggests that this looping may constitute a feedback mechanism to repress *TERT* expression in cells with long telomeres, a phenomenon termed telomere position effect (TPE)^38–40^. We explored a possible TPE in neuroblastoma cells with long telomeres and quantified the abundance of chromatin in repressive states 5Mb from chromosomal ends and performed a similar analysis genome-wide to verify that any change at chromosomal ends do not represent a genome-wide difference in repressive marks among these cells. We focused on the three repressive states: ChromHMM 15 (polycomb repressed, enriched for H3K27me3), ChromHMM 16 (heterochromatin, enriched for H3K9me3), and ChromHMM 17 (repressed, enriched for both H3K9me3 and H3K27me3). We found that, relative to the whole genome, repressive marks are more abundant near chromosomal ends in all neuroblastoma cells. However, chromosomal termini from neuroblastoma cells with long telomeres are enriched for repressive marks relative to those from short telomere/telomerase-positive neuroblastoma cells (Figure 5A, Supplementary Figure S4, Supplementary Table 1A-D). Additionally, the majority of *TERT*-positive neuroblastoma cells are *MYCN*-amplified. This suggests that the enrichment of repressive marks at chromosomal ends in cells with long telomeres relative to those with short telomeres may represent a reduction in repressive marks due to *MYCN* amplification rather than TPE. To test this possibility, we induced *MYCN* expression in a neuroblastoma cell line with long telomeres that has a doxycycline inducible *MYCN* transgene, SKNMM^MYCN^^16^. We found no significant change in the abundance of the repressive ChromHMM states at chromosomal ends in this cell line upon *MYCN* induction (Supplementary Figure S4). Taken together, these data suggest that long telomeres are associated with repressive chromatins at the chromosomal ends in neuroblastoma cells.

**Figure 5:**
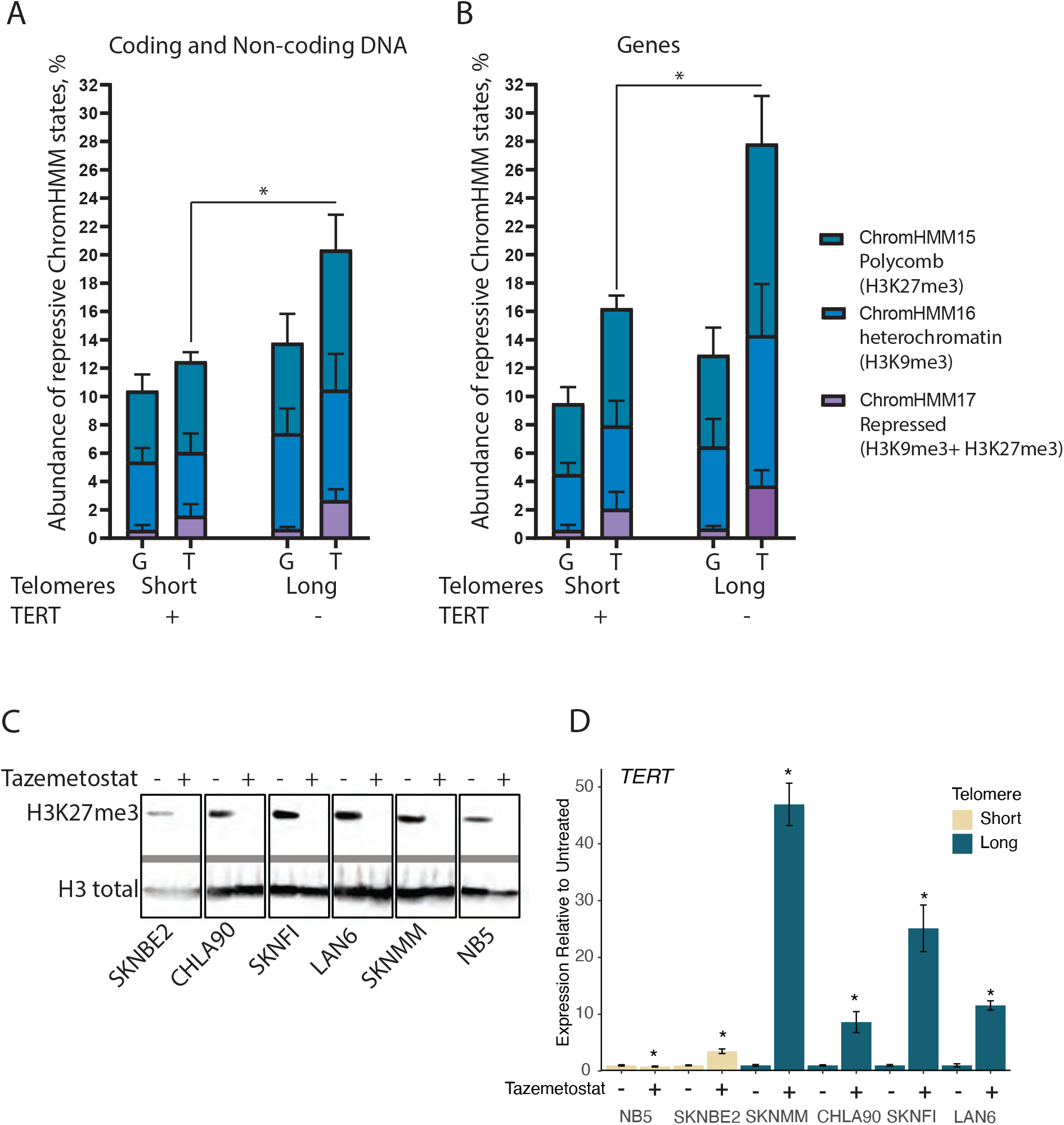
A-B; Telomere position effect (TPE) suppresses genes at chromosomal ends of neuroblastoma cells with long telomeres. The average of the percentage of repressive chromatin ChromHMM states at chromosomal ends, subtelomeric region (T) and genome-wide (G) for all coding and non-coding sequences (A) and genic regions only (B) of neuroblastoma cells with short and long telomeres. **C-D; Pharmacological inhibition of EZH2 induces *TERT* expression in *TERT*-negative neuroblastoma cells**. **C-** Western Blot for H3K27me3 and H3K27 total in neuroblastoma cells after treatment with tazemetostat or the drug vehicle only. **D**-Fold change of *TERT* expression in the same cells after the same treatment measured by qRT-PCR. * indicates p< 0.05, error bars represent standard deviation

It was previously shown that TPE represses genes at long distances from chromosomal ends^38–40^. To test if long telomeres have differential repressive effects on genes at chromosomal ends, we further analyzed chromatin repressive marks at chromosomal ends in genic (genes + 2Kb upstream and downstream) regions. Again, repressive marks were more abundant at genes located near the chromosomal ends relative to whole genome in all neuroblastoma cells. However, this enrichment was more evident in neuroblastoma cells with long telomeres (Figure 5B, Supplementary Figure S4, Supplementary table 1A-D). *MYCN* induction in SKNMM^MYCN^ cells did not significantly alter this enrichment (Supplementary Figure S4). These data suggest that TPE is detected in neuroblastoma cells with long telomeres and TPE may also play a role in repressing *TERT* expression.

### Inhibition of EZH2 induces TERT expression in neuroblastoma cells with long telomeres

EZH2 is the enzymatically active subunit of the polycomb repressive complex 2 (PRC2), methylating lysine 27 on histone H3 tails to repress gene transcription^22^. To examine the impact of H3K27me3 in the transcriptional silencing of the *TERT* locus in neuroblastoma, we treated neuroblastoma cell lines with long telomeres (SKNFI, SKNMM, CHLA-90 and LAN6) and *MYCN*-amplified neuroblastoma cell lines with short telomeres (SKNBE2 and NB5) with the FDA-approved EZH2 inhibitor, Tazemetostat, and measured *TERT* expression. We treated neuroblastoma cell lines with a range of Tazemetostat doses (0-20 μM) and at two time points (10 days and 21 days) and found treating cells with 20 μM of Tazemetostat for 21 days reduced methylated H3K27 below the limit of detection as assessed by Western blot (Figure 5C). Following 21 days of treatment with 20 μM of Tazemetostat, we observed an induction in *GREB1* and *SPTBN2* (Supplementary Figure S5 A & B), genes previously shown to be transcriptionally repressed by H3K27me3 in neuroblastoma cells^33^. Interestingly, we also observed that inhibiting EZH2 de-repressed *TERT* and induced expression in neuroblastoma cells with long telomeres (Figure 5D, Supplementary Figure S5C). However, this activation of *TERT* expression was not accompanied by changes in C-circle levels, a canonical ALT-associated hallmark (Supplementary Figure S5D). The enrichment of H3K27me3 at the *TERT* proximal promoter in neuroblastoma cells with long telomeres, and the induction of *TERT* expression following Tazemetostat treatment in those cell lines, support the notion that PRC2 plays an important role in repressing telomerase in telomerase-independent neuroblastoma.

## Discussion

The epigenetic control of *TERT* expression is not well understood^28^. Here, we leveraged our extensive neuroblastoma epigenetic database^16^ to understand how the *TERT* promoter is regulated in neuroblastoma. The *TERT* proximal promoter spanning nucleotides +51 to −650 from the TSS is composed of a core promoter and an upstream region that has been previously described as the *TERT* hypermethylated oncological region (THOR)^17,30^. Several transcriptional factors, including MYC family members, bind to the core promoter^17^. Indeed, we have shown that MYCN binds to *TERT* core promoter in *MYCN*-amplified cells. Broadly, cancer cells and immortalized cell lines are hypermethylated at CpGs in the core *TERT* promoter and THOR region^27^, including cancers of the central nervous system^41^. However, we show that neuroblastoma cells are exceptional, displaying CpG hypomethylation within the core promoter and THOR, similar to stem cells and non-malignant somatic cells^27, 41^. Notably, the developing adrenal medullary cells show a similar CpG hypomethylation at *TERT* promoter DNA. Taken together, these data are consistent with the idea that most neuroblastoma tumors arise from sympathetic progenitor cells arrested during development.

The role of *TERT* promoter DNA methylation in the regulation of *TERT* expression is not well-established^28,42^. Generally, CpG methylation is considered a repressive mark, blocking the binding of transcription factors to promoters^22^. Paradoxically, *TERT* expression is associated with promoter DNA hypermethylation^42^. While it has been suggested that *TERT* promoter DNA methylation prevents CTCF from binding and repressing *TERT* expression^43^, several studies have shown that monoallelic *TERT* expression preferentially comes from alleles with hypomethylated promoters^26,27,44,45^. This *TERT* promoter CpG hypermethylation phenomenon may be attributed as an attempt by cancer cells to sustain only the minimal level of telomerase needed for telomere length maintenance^27^. This hypothesis is supported by the fact that *TERT* amplification is relatively rare in cancer and most telomerase-positive cancer cells have short telomeres^2,7^.

To gain a more comprehensive appreciation of the epigenetic regulation of the *TERT* proximal promoter, we complemented our CpG methylation data with extensive chromatin analysis. *TERT* is a known target of MYCN^*17*^, and consistent with this, our data showed that *MYCN*-amplified neuroblastoma cells express telomerase. Additionally, we found that the NBLS cell line, which does not have a *MYCN* amplification, but does have moderate MYCN expression^46^, also expresses telomerase. The *TERT* promoter in telomerase-positive neuroblastoma cell lines showed enrichment for both the repressive mark H3K27me3 and the active mark H3K4me3. While this combination of repressive and active marks has been observed in other regulated genes and described as a bivalent promoter^37^, it is also possible that these marks do not co-exist on the same allele. Indeed, previous studies have shown monoallelic *TERT* expression^26,44^.

In addition to telomerase-dependent neuroblastoma cells, a subset of high-risk neuroblastoma cells is telomerase-independent. While neuroblastoma cells harboring *ATRX* mutations are thought to rely on ALT to maintain telomere lengths^18^, a recent study has shown that neuroblastoma tumors can also be negative for both ALT and telomerase. These tumors are characterized by very long telomeres (>20 kbps) that allow these cells to pass through many replication cycles without inducing senescence and apoptosis^6^. We found that neuroblastoma cells with long telomeres, with little to no telomerase expression, possess a unique epigenetic signature. Strikingly, neuroblastomas with long telomeres display DNA CpG hypomethylation and enrichment of the chromatin repressive mark H3K27me3 that extends for tens of kilobases around the *TERT* locus and includes neighboring genes. Previous work has shown a similar signature of H3K27 trimethylation and DNA hypomethylation (termed DNA methylation valleys) in some developmentally repressed genes^47^. Interestingly, similar to *TERT* in cancer, activation of genes in DNA methylation valleys is associated with DNA hypermethylation^47^. Overall, these data suggest that neuroblastoma with long telomeres, including ALT-positive tumors, may arise from sympathetic progenitor cells at a developmental stage different from telomerase-dependent neuroblastomas in which the *TERT* locus is silenced by PRC2 in a DNA methylation valley. Indeed, it has been shown that the *TERT* promoter will undergo changes in epigenetic marks during development^48^, although not specifically in developing sympathetic ganglion cells. Characterizing the *TERT* epigenetic signature of sympathetic neuron progenitor cells at different developmental stages may provide some key insights in different telomere maintenance mechanisms employed by neuroblastoma.

In this study, we showed enrichment of chromatin repressive marks close to chromosomal ends relative to whole genome. These repressive marks are more evident at genic rather than intergenic regions. Notably, cells with long telomeres had more abundant repressive marks and repressive states relative to telomerase-positive neuroblastoma cells (Figure 5A-B, Supplementary Figure S4, Supplementary Table 1A-D), suggesting a TPE. The telomere position effect was previously described as a negative feedback mechanism to inhibit *TERT* expression in cells with long telomeres^38,39^; *TERT* is close to the chromosome 5p end. However, this observation does not explain the repression of other genes in the region. An intriguing idea is that repression of subtelomeric genes in cells with long telomeres may work as a barrier to acquire long telomeres. In this model, cells with long telomeres, for example ALT positive cells, must find a way to compensate for the repression of genes at chromosomal ends. TPE may also explain the relatively stable short telomeres detected in most telomerase positive cancer cells^38^. Minimal telomerase expression will balance the positive selection for continual cellular proliferation without going through telomere attrition-inducing senescence and a potential negative selection of suppressing the expression of subtelomeric genes. Thus, future studies are needed to explore these possibilities.

In conclusion, we show that the *TERT* region is differentially regulated in neuroblastoma cells depending on the mechanism of telomere maintenance. In addition, our data demonstrate that promoter DNA methylation is unique in neuroblastoma, with a CpG methylation pattern similar to somatic and stem cells, supporting the hypothesis that neuroblastoma represents an arrested differentiation of sympathetic neuron progenitors. Overall, these findings reveal new insights in the epigenetic regulation of the *TERT* promoter in neuroblastoma. Notably, in terms of treatment, the utility of EZH2 inhibitors in neuroblastoma may in part be influenced by telomere maintenance mechanism.

## Material and methods

### Cell lines and culture

U2OS, 293T, SKNBE2, IMR32 and SKNFI cells were obtained from the American Type Culture Collection (ATCC, Manassas, VA). CHLA90 and LAN6 cells were obtained from the Children Oncology Group (COG). NB-5 cells were available at St. Jude Children’s Research Hospital. SKNMM cells were a generous gift from Dr. Nai-Kong Cheung from the Memorial Sloan Kettering Cancer Center while NBLS cells were a kindly provided by Dr. Garrett M Brodeur from the Children Hospital of Philadelphia (CHOP). 293T, SKNFI, NB5 and SKNFI cells were grown in DMEM medium (Lonza, Cat# 12-614F) supplemented with 10% FBS and 1X Penicillin/Streptomycin /L-Glutamine (Gibco, Cat# 1037-016). NBLS, SKNMM and LAN6 cells were maintained in RPMI-1640 medium (Lonza, Cat# 12-167Q) supplemented with 1X Penicillin/Streptomycin /L-Glutamine (Gibco, Cat# 1037-016) and 10% FBS. IMR-32 cells were grown in EMEM cells supplemented by 1X Penicillin/Streptomycin (ATCC, Cat# 30-2003) and 10% FBS while U2OS cells were grown in McCoy’s 5A medium (Sigma, Cat# M8403) supplemented with 1X Penicillin/Streptomycin (Gibco, Cat# 15140-122) and 10% FBS. The cells were checked for mycoplasma contamination using Universal Mycoplasma Detection Kit (ATCC, Cat. # 30-1012K) following the manufacturer’s protocol and the identity of the cell lines were confirmed periodically using short tandem repeat (STR) profiling.

### Fluorescent In Situ Hybridization (FISH) for Telomeres and MYCN

FISH was done as previously described^16^. In brief, telomere interphase FISH was performed using fluorescein isothiocyanate-labeled PNA telomere probe (DAKO, Cat. # 5327) following the manufacturer’s protocol. For *MYCN* FISH, *NMYC* BAC DNA (*RP11-1183P10*) and human chromosome 2 control (2q11.2) BAC DNA (*RP11-527J8*) were labelled with red-dUTP and green-dUTP, respectively, using nick translation kit (Molecular Probes, Cat# AF594). The samples were denatured 90 °C for 12 min. Hybridization with the labelled probes mixed with sheard human DNA was done overnight at 37°C on a ThermoBrite (Abbott, Chicago, IL) in the hybridization solution (50% formamide, 10% dextran sulfate, and 2× SSC solution). Excess probes were washed and 4,6-diamidino-2-phenylindole (DAPI) (1 μg/ml) was used as a counter stain. Image capture and analysis was done using Zeiss Axio Imager.Z2 microscope and GenASIs scanner (Zeiss, Thornwood, NY).

### Telomere quantitative polymerase chain reaction (qPCR)

Quantification of telomeric repeats was performed using qPCR as previously described^16^. In brief, DNA was extracted using DNAeasy Kit (Qiagen, Cat. # 69504) following the manufacturer’s instructions. qPCR reaction was done in 10-μl total reaction volume; containing 10-ng DNA template, 10-mM primer mix and 5-μl SYBR green Select master mix CFX (Thermo Fisher, Cat. #4472942). Every reaction was done in triplicate and for every sample, 2 sets of primers were used; one for the telomeric sequence and the other for the control locus *RPLP0*. Primer sequences are listed in the supplementary table S2. The average C(t) value of the telomeric repeat reactions was subtracted from the average C(t) value of the control locus reactions to get ΔC(t). Quantification of the telomeric repeats were presented as fold changes relative to telomerase positive 293T cells or to the untreated condition.

### Real-time quantitative polymerase chain reaction (RT-qPCR)

RNA was extracted using RNeasy kit (Qiagen, Cat # 74004) following the manufacturer’s protocol. Extracted RNA was reverse transcribed using the iScript Advanced cDNA synthesis kit (Bio-Rad, Cat# 1725038). RT-qPCR was performed on the equivalent of 50 ng of cDNA with 500 nM of primer using SsoAdvanced Universal SYBR green Supermix (Bio-Rad, Cat# 1725271). The following conditions were used for qPCR: 98 °C for 30 s and 35 cycles at 98 °C for 15 seconds and 60 °C for 30 seconds. The expression of *TERT, TERC, GREB1*, and *SPTBN2* were normalized to *HPRT* (Supplemental Table S2). Results were analyzed using CFX Manager Software 3.1 (Bio-Rad) to assess the relative gene expression levels of normalized target genes.

### C-Circle assay

We performed the ALT-associated C-circle assay following a previously described protocol with some modifications^49^. Briefly, DNA was extracted from the cells using DNeasy Blood and Tissue Kit (Qiagen, Cat# 69506) protocol and then was purified using QiaQuick PCR Purification Kit (Qiagen, Cat # 28104) following the manufacturer’s protocols. Rolling sample amplification reaction mix (containing: 0.2mg/ml bovine serum albumin (BSA), 0.1% Tween-20, 1X Phi29 reaction buffer and 1mM of dGTP, dATP and dTTP) was added to 150-ng DNA from each sample with or without 2 units of Phi29 polymerase (New England Biolabs, Cat# M0269L) in 20-μl total reaction volume and incubated at 30 °C for 8 hours. The reaction was then inactivated by incubating samples for 20 minutes at 65°C. Reactions were blotted onto positively-charged nylon membrane (Roche) and cross linked using Stratagene UV Stratalinker (Stratagene, San Diego, CA). The membrane was washed with 2X saline sodium citrate (2XSCC) buffer with 0.1% SDS, blocked using DIG Easy-Hyb for 1 hour at 42°C (Roche, Cat#11093274910). Telomeric repeats were probed by incubating the membrane with telomeric probe containing digoxigenin (5’CCCTAACCCTAACCCTAACCCTAA-DIG; Integrated DNA Technologies, Coralville, IA; Table S1) overnight at 42°C DIG Easy Hyb™ buffer (Roche). Blots were washed in 2X saline-sodium citrate (SSC) buffer with 0.1% SDS at room temperature (RT), followed by 0.2X SSC 0.1% SDS at 50°C, and blocked in 5% milk TBST for 30 minutes at RT. Blots were incubated in anti-Digoxigenin-AP, Fab fragments diluted 1:10,000 in the blocking buffer for 30 minutes at room temperature (Roche, Cat#11093274910) following the manufacturer’s protocols. Blots were washed in 2X SSC buffer with 0.1% SDS at RT. C-circle signal was detected using the CDP-Star kit (Roche) according to the manufacturer’s instructions and imaged using the ChemiDoc™ Imaging System (BioRad, Hercules, CA). U2OS cells (serial dilution from 0-50 ng DNA) were used as an ALT positive control while PC3 cells (150-ng DNA) were used as a negative control.

### Western Blot

The cells were harvested from a 100-mm tissue culture plate by removing the medium, washing twice with PBS (pH 7.4) and scraping in 1-ml PBS. The cells were then pelleted by centrifugation at 13,000 rpm for 1 min at 4 °C. Cells were resuspended in 250-μl Triton extraction buffer containing 0.5% Triton-X 100, Halt protease inhibitor to a final concentration of 1X (Thermo Fischer, Cat. #7834) and ethylenediaminetetraacetic acid (EDTA) to a final concentration of 5 mM. Cells were incubated in the lysis buffer for 10 minutes on a horizontal shaker at 4 °C followed by centrifugation at 6,500 x g for 10 minutes at 4 °C to collect the nuclei. The supernatant was discarded, and the nuclei were washed with 125-μl of ice-cold Triton extraction buffer. The samples were then centrifuged again 6,500 x g for 10 minutes at 4 °C and the supernatant was discarded. The pelleted nuclei were re-suspended in 100-ml of 0.2N HCl containing 10% glycerol and incubated on a gentle horizontal shaker at 4°C overnight. The debris were pelleted and discarded by centrifuging the tubes at 14,000X for 10 minutes at 4 °C. 90-μl of the supernatant was transferred to a new tube containing 10-μl neutralization buffer containing: 2M NaOH, 1X Halt protease inhibitor (Thermo Fischer, Cat. #7834) and 10mM DTT. Protein concentration was determined using BCA protein assay kit (Thermo Fischer, Cat. #232225), following manufacturer’s instructions. Equal amount of protein from different samples and 5-μl of Odyssey One-Color Protein Molecular Weight Marker (LiCore Cat# 928-40000) were resolved on 4—15% sodium dodecyl sulfate polyacrylamide gel (SDS-PAGE, Biorad Cat#4561086) then the protein was transferred to a nitrocellulose membrane at 4 °C overnight at 30 volts. The membrane was washed and blocked for 1 hour at room temperature using Odyssey Blocking buffer, PBS (LiCore Cat# 927-40000). The membranes were probed with Anti-histone H3 Rabbit Ab (Cell Signaling, Cat# 9715S) or anti-tri-methyl Histone H3 (H3K27me3, Cell Signaling, Cat# 9733S) diluted 1:1000 for 1 hour at room temperature then washed three times, 5 minutes each with PBS supplemented with 0.1% Tween-20. The membranes then were incubated with IRDye 680CW goat anti-rabbit antibodies (1:5000, LiCore, 925-32211) were used for 1 hour at room temperature followed by washing for three times, 5 minutes each, with PBS supplemented with 0.1% Tween-20. Odyssey CLX infrared gel imaging system (Li-Cor Biosciences, Lincoln, NE) was used for scanning the membranes.

---

**Supplementary figure S1:**
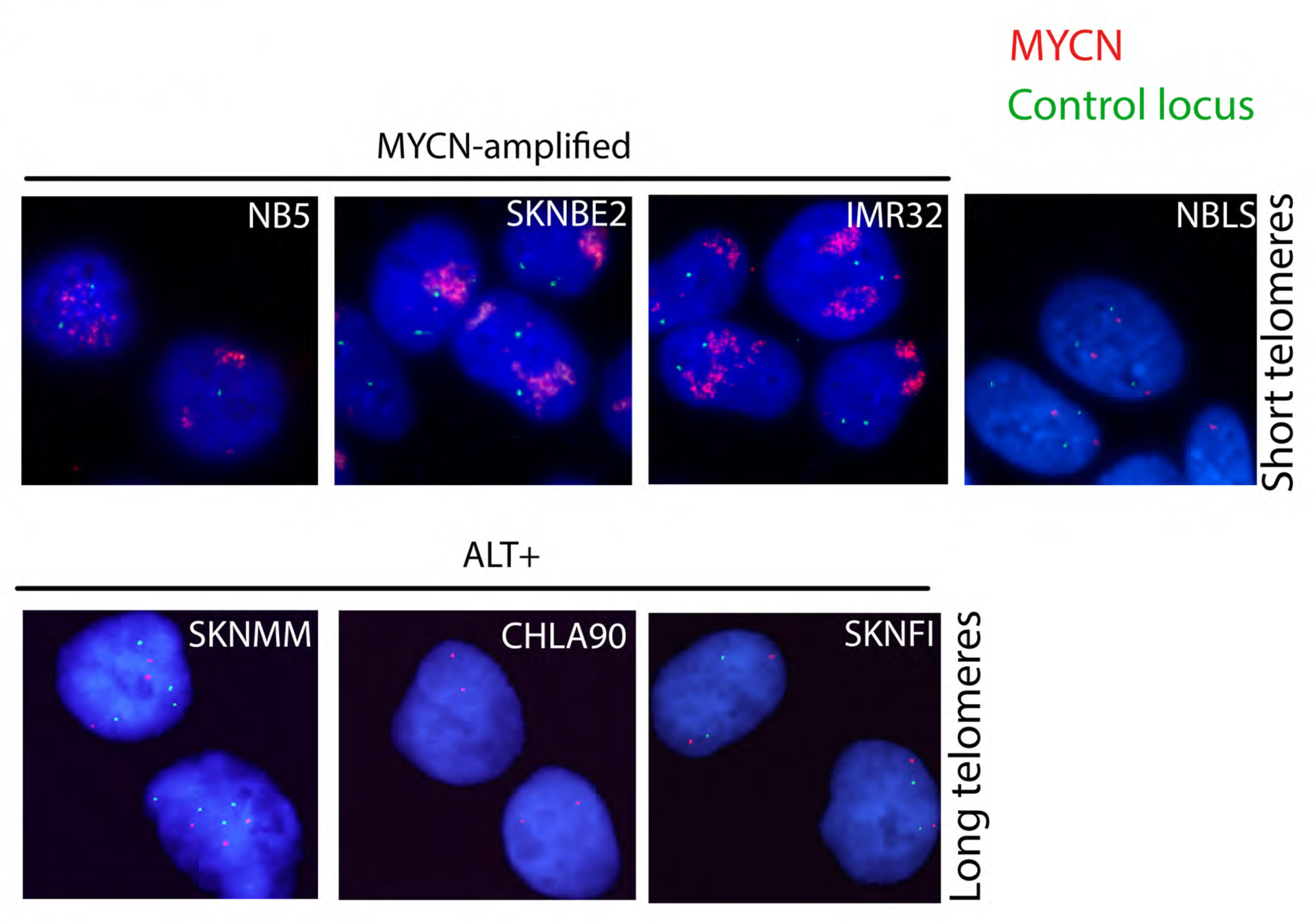
Representative fluorescence microscopy images of MYCN-FISH (red) and a control locus (green) of seven neuroblastoma cell lines used in this study.

**Supplementary figure S2:**
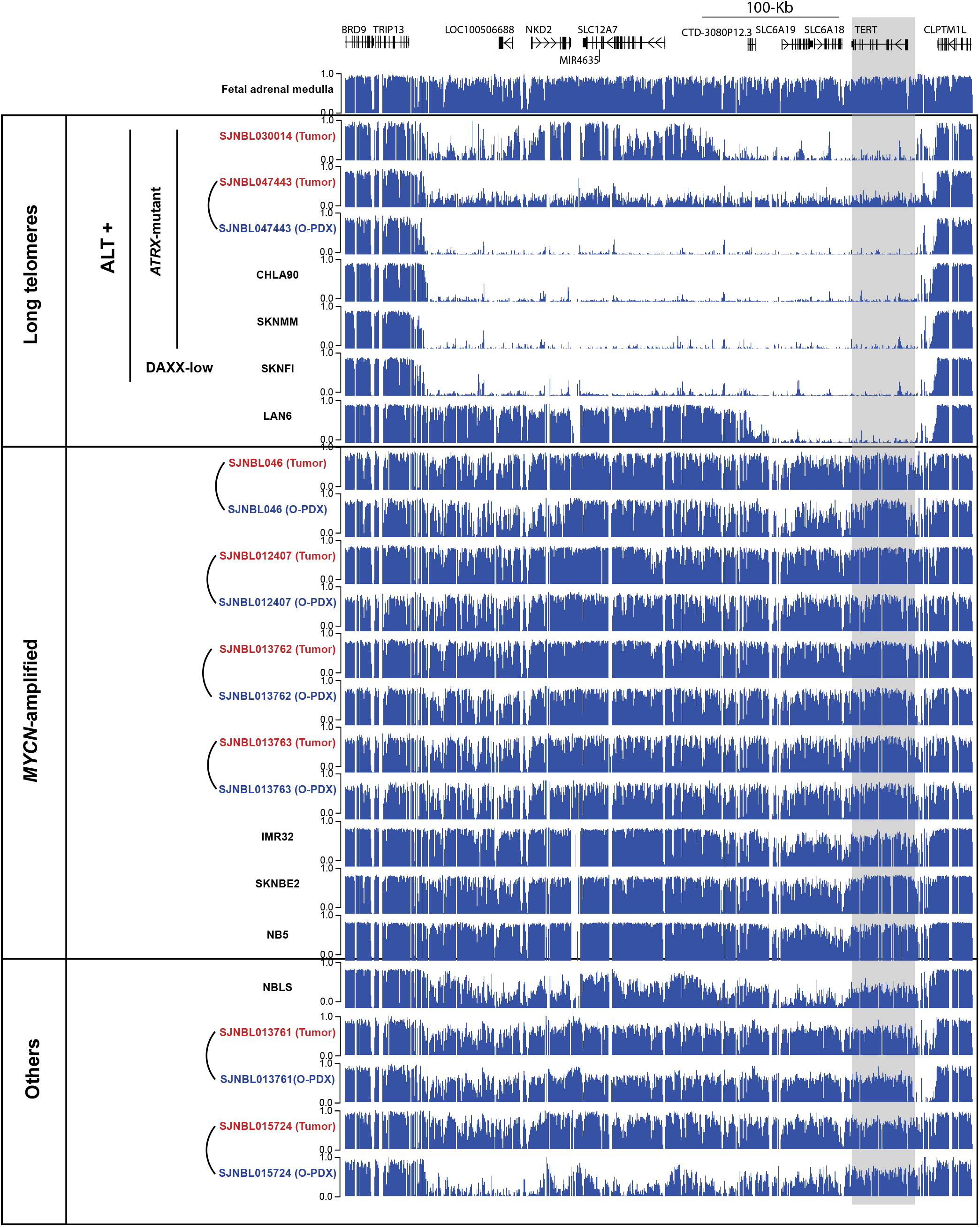
The DNA hypomethylated region extends for tens of thousands of kilobases in neuroblastoma cell cells with long telomeres. *TERT* locus is shaded in grey.

**Supplementary figure S3:**
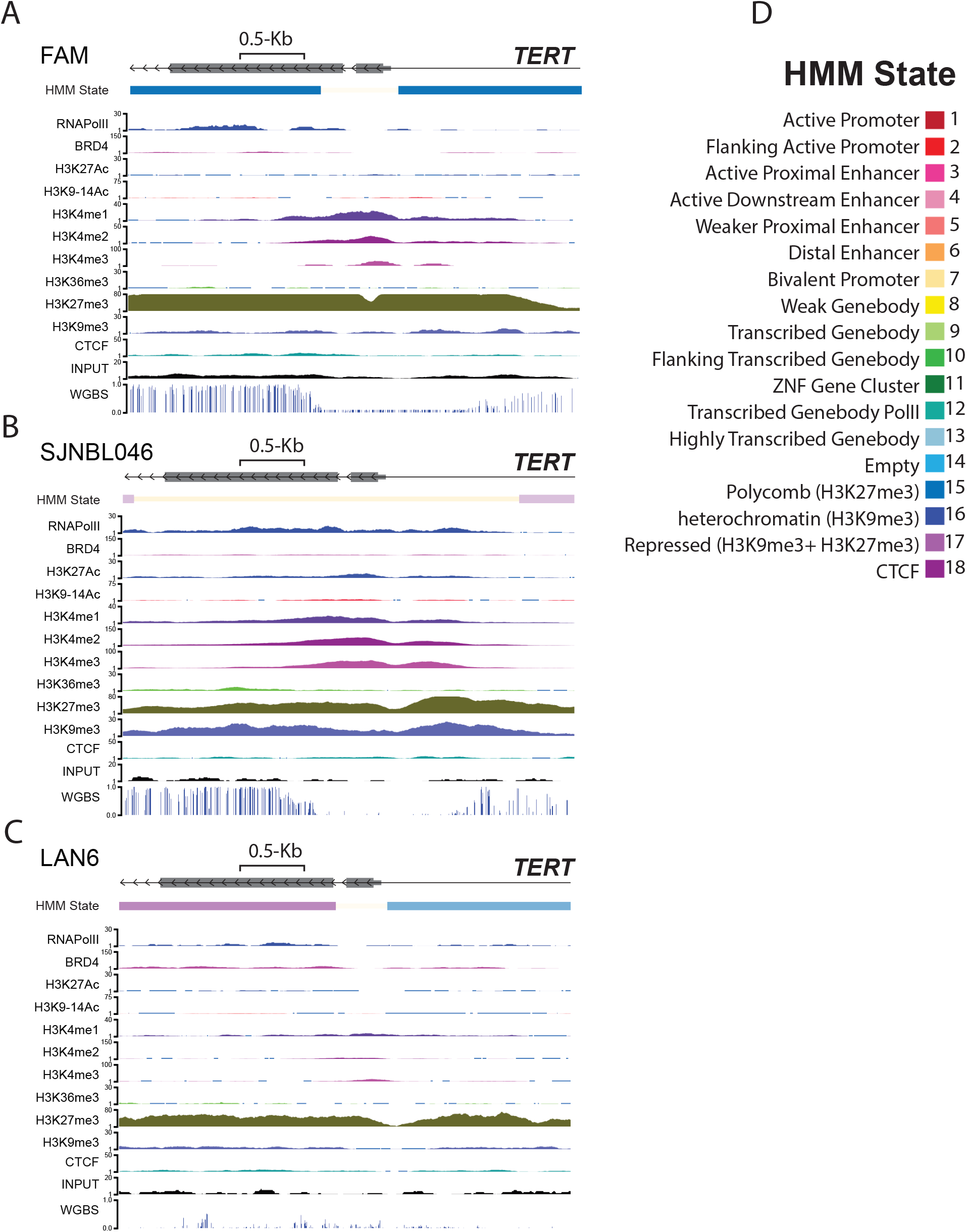
ChromHMM state, all ChIP-Seq tracks and whole geneome bisulfite sequencing (WGBS) for fetal adrenal medulla (FAM, **A**), *MYCN*-amplified O-PDX, SJNLB046 (**B**) and the neuroblastoma cell line with ever-shorter telomere, LAN6 (C). The color codes for different ChromHMM states are shown in **D.**

**Supplementary figure S4:**
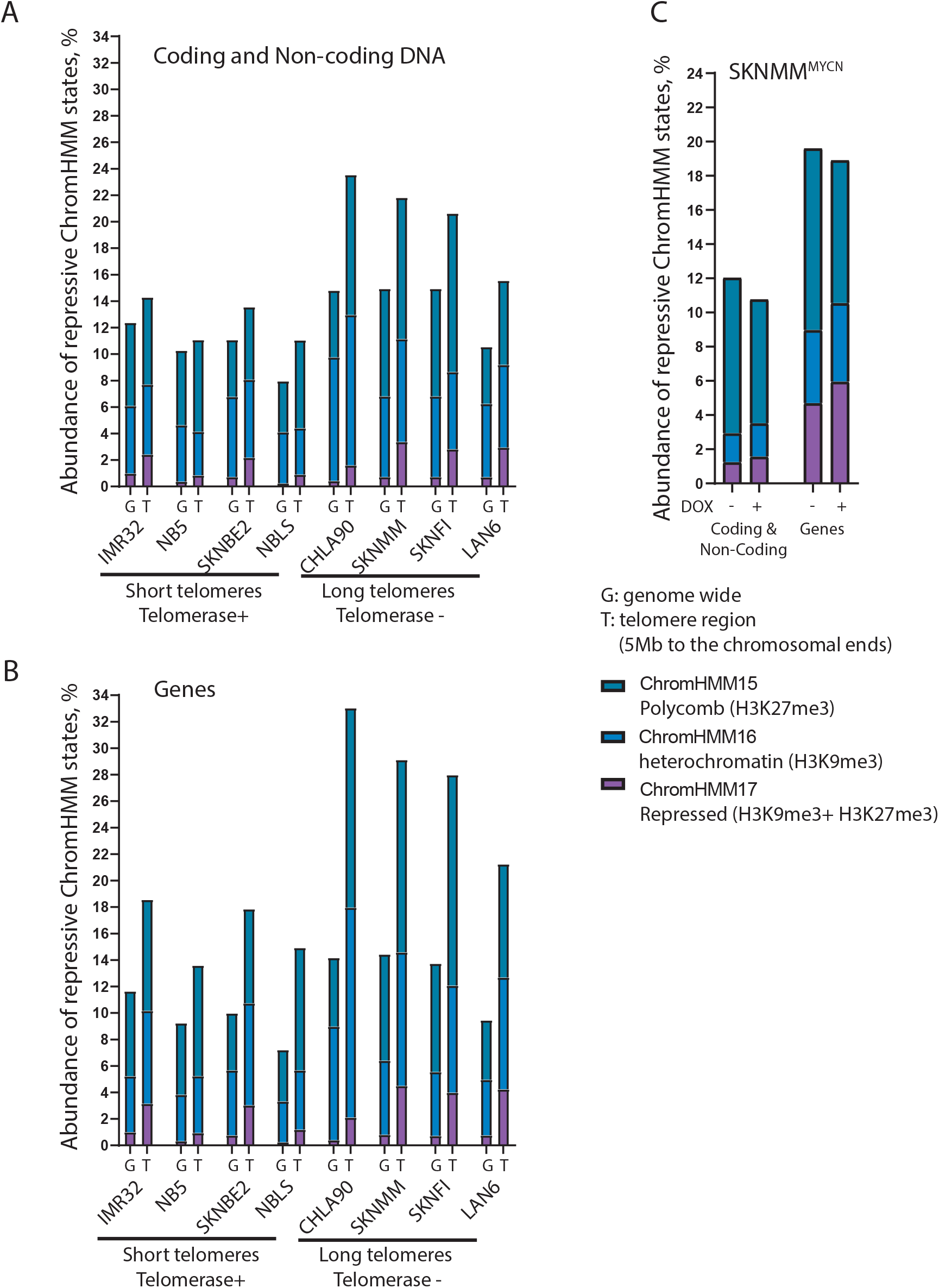
Abundance of repressive chromatin ChromHMM states of 5 Mb at the chromosomal ends, subtelomeric region (T) and genome-wide (G) in neuroblastoma cells with short versus long telomeres, the whole coding and non-coding region (A) and the genic region (B). Repressive chromatin states at the chromosomal ends of the neuroblastoma cells with long telomeres with or without induction of MYCN transgene (C).

**Supplementary figure S5:**
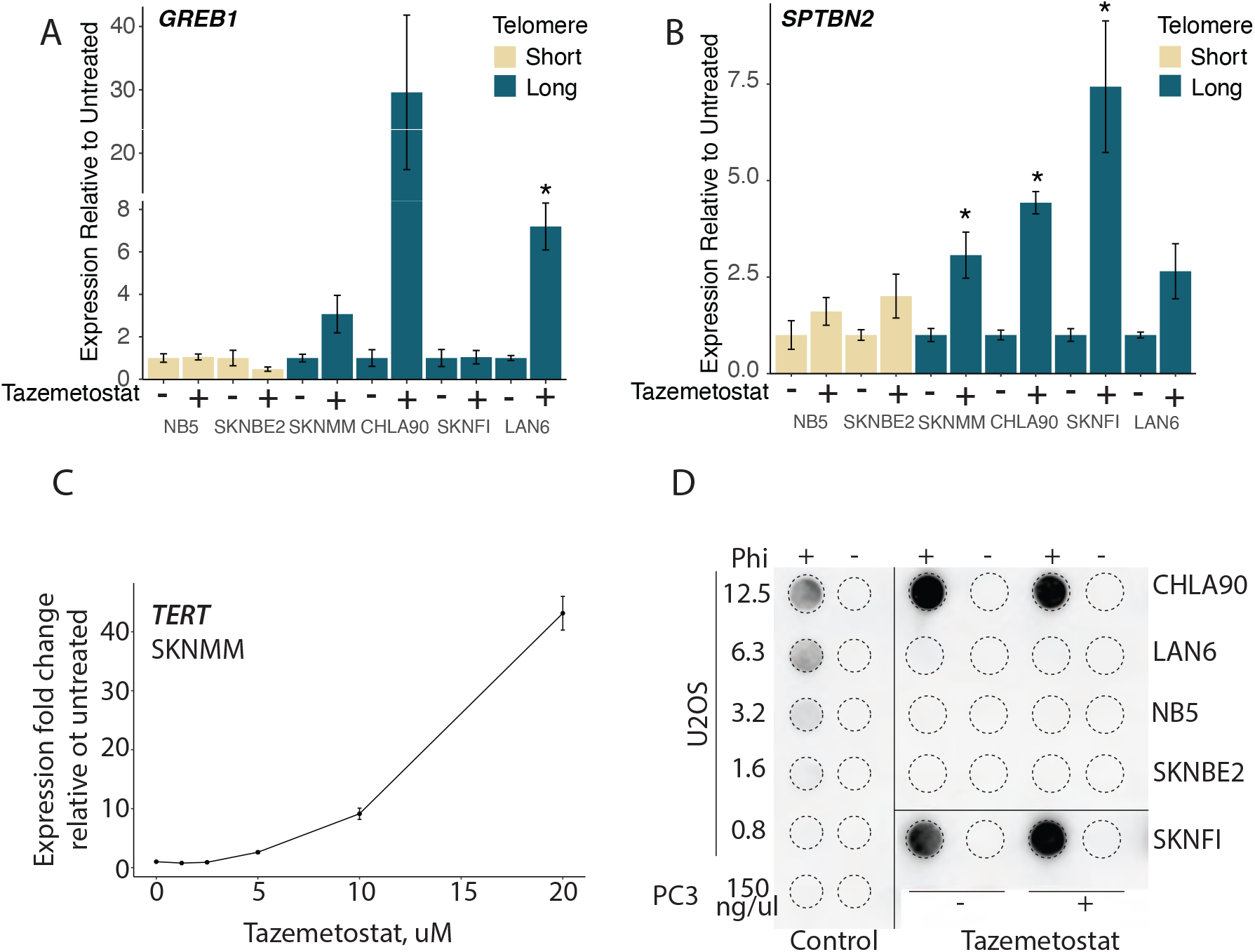
Treatment of neuroblastoma cell lines with Tazemetostat or the drug vehicle only, **A)** Fold change measured by RT-qPCR for **A)** *GREB1* and **B)** *SPTBN2*. * Indicates p< 0.05, error bars represent standard deviation. **C)** Fold change of *TERT* expression of SKNMM cells treated with different concentration of EZH2 inhibitor Tazemetostat for 3 weeks as measured using RT-qPCR. **C)** C-circle assay blot with U2OS and PC3 cells used as positive and negative controls, respectively. The amount of DNA in nanograms (ng) is shown on the blot. All reactions were done in the presence of absence of the Phi 29 (Phi) DNA polymerase.

**Supplementary table1A:** abundance of different ChromHMM in neuroblastoma cell lines. Genes including 2Kb up and downstreams in the 5 Mb of subtelomeric sequences

|  | Short telomeres<br>Telomerase + |  |  |  | Long telomeres<br>Telomerase - |  |  |  |
| --- | --- | --- | --- | --- | --- | --- | --- | --- |
|  | IMR32 | NB5 | SKNBE2 | NBLS | CHLA90 | SKNMM | SKNFI | LAN6 |
| State1 | 1.31 | 0.54 | 1.24 | 1.46 | 1.18 | 1.42 | 1.22 | 1.42 |
| State2 | 0.04 | 0.01 | 0.14 | 0.03 | 0.05 | 0.14 | 0.04 | 0.22 |
| State3 | 1.13 | 0.88 | 2.31 | 1.06 | 0.66 | 0.77 | 0.63 | 2.10 |
| State4 | 1.01 | 1.41 | 2.84 | 1.89 | 0.47 | 1.05 | 0.75 | 2.34 |
| State5 | 1.94 | 3.36 | 1.40 | 2.32 | 2.13 | 1.81 | 1.26 | 1.36 |
| State6 | 4.52 | 4.94 | 7.23 | 3.84 | 1.48 | 2.01 | 1.84 | 4.83 |
| State7 | 0.45 | 0.65 | 1.04 | 0.25 | 0.56 | 1.05 | 0.49 | 0.60 |
| State8 | 11.32 | 12.74 | 8.91 | 13.05 | 13.48 | 12.03 | 11.85 | 9.20 |
| State9 | 7.43 | 5.95 | 3.65 | 6.06 | 2.83 | 8.27 | 8.56 | 6.16 |
| State10 | 1.07 | 0.69 | 1.94 | 0.56 | 0.27 | 1.28 | 0.34 | 2.01 |
| State11 | 2.08 | 1.31 | 1.93 | 1.09 | 0.47 | 0.84 | 0.96 | 1.41 |
| State12 | 2.26 | 1.48 | 1.83 | 2.70 | 2.28 | 1.94 | 3.18 | 1.91 |
| State13 | 0.09 | 0.08 | 0.18 | 0.10 | 0.04 | 0.28 | 0.04 | 0.26 |
| State14 | 46.13 | 51.53 | 47.17 | 50.37 | 40.50 | 37.49 | 40.50 | 44.24 |
| State15 | 8.38 | 8.36 | 7.11 | 9.26 | 15.06 | 14.54 | 15.90 | 8.55 |
| State16 | 7.02 | 4.29 | 7.70 | 4.48 | 15.86 | 10.07 | 8.09 | 8.45 |
| State17 | 3.16 | 0.94 | 3.04 | 1.19 | 2.11 | 4.51 | 3.99 | 4.24 |
| State18 | 0.67 | 0.87 | 0.35 | 0.30 | 0.56 | 0.49 | 0.36 | 0.71 |

**Supplementary table 1B:** abundance of different ChromHMM in neuroblastoma cell lines. Genic and intergenic regions at 5Mb of subtelomeric sequences.

|  | Short telomeres<br>Telomerase + |  |  |  | Long telomeres<br>Telomerase - |  |  |  |
| --- | --- | --- | --- | --- | --- | --- | --- | --- |
|  | IMR32 | NB5 | SKNBE2 | NBLS | CHLA90 | SKNMM | SKNFI | LAN6 |
| State1 | 0.68 | 0.28 | 0.64 | 0.79 | 0.63 | 0.74 | 0.64 | 0.75 |
| State2 | 0.02 | 0.00 | 0.07 | 0.01 | 0.03 | 0.08 | 0.02 | 0.12 |
| State3 | 0.71 | 0.52 | 1.30 | 0.63 | 0.40 | 0.46 | 0.38 | 1.25 |
| State4 | 0.69 | 0.86 | 1.72 | 1.20 | 0.28 | 0.64 | 0.47 | 1.51 |
| State5 | 1.08 | 1.91 | 0.77 | 1.34 | 1.21 | 1.02 | 0.71 | 0.75 |
| State6 | 2.87 | 3.07 | 4.54 | 2.49 | 0.89 | 1.28 | 1.15 | 3.17 |
| State7 | 0.27 | 0.40 | 0.62 | 0.16 | 0.37 | 0.67 | 0.31 | 0.35 |
| State8 | 5.83 | 6.58 | 4.60 | 6.94 | 7.11 | 6.17 | 6.11 | 4.81 |
| State9 | 3.76 | 3.03 | 1.86 | 3.19 | 1.48 | 4.20 | 4.37 | 3.20 |
| State10 | 0.54 | 0.35 | 0.98 | 0.29 | 0.14 | 0.66 | 0.18 | 1.05 |
| State11 | 1.19 | 0.69 | 1.08 | 0.61 | 0.26 | 0.46 | 0.51 | 0.78 |
| State12 | 1.23 | 0.80 | 0.99 | 1.50 | 1.27 | 1.07 | 1.71 | 1.06 |
| State13 | 0.04 | 0.04 | 0.09 | 0.05 | 0.02 | 0.14 | 0.02 | 0.13 |
| State14 | 66.35 | 69.78 | 66.95 | 69.53 | 62.00 | 60.27 | 62.57 | 65.03 |
| State15 | 6.57 | 6.94 | 5.47 | 6.64 | 10.58 | 10.66 | 11.97 | 6.35 |
| State16 | 5.29 | 3.29 | 5.90 | 3.49 | 11.36 | 7.77 | 5.82 | 6.24 |
| State17 | 2.42 | 0.85 | 2.18 | 0.91 | 1.60 | 3.38 | 2.83 | 2.96 |
| State18 | 0.45 | 0.60 | 0.24 | 0.22 | 0.37 | 0.33 | 0.24 | 0.49 |

**Supplementary table1C:** abundance of different ChromHMM in neuroblastoma cell lines. Genes including 2Kb up and downstream, genome-wide.

|  | Short telomeres<br>Telomerase + |  |  |  | Long telomeres<br>Telomerase - |  |  |  |
| --- | --- | --- | --- | --- | --- | --- | --- | --- |
|  | IMR32 | NB5 | SKNBE2 | NBLS | CHLA90 | SKNMM | SKNFI | LAN6 |
| State1 | 0.77 | 0.37 | 0.69 | 0.83 | 0.84 | 0.90 | 0.71 | 0.84 |
| State2 | 0.01 | 0.00 | 0.03 | 0.01 | 0.02 | 0.04 | 0.01 | 0.05 |
| State3 | 0.58 | 0.68 | 0.87 | 0.44 | 0.64 | 0.59 | 0.60 | 0.89 |
| State4 | 0.99 | 1.40 | 1.90 | 1.33 | 0.72 | 1.33 | 1.17 | 1.51 |
| State5 | 1.27 | 2.23 | 1.05 | 1.62 | 1.61 | 1.68 | 1.05 | 1.18 |
| State6 | 2.10 | 4.01 | 2.96 | 2.81 | 2.04 | 2.53 | 1.84 | 3.00 |
| State7 | 0.19 | 0.33 | 0.37 | 0.12 | 0.25 | 0.42 | 0.25 | 0.21 |
| State8 | 9.74 | 10.29 | 8.01 | 9.97 | 10.12 | 11.57 | 9.94 | 8.77 |
| State9 | 4.51 | 3.03 | 2.01 | 2.37 | 0.96 | 3.57 | 4.99 | 3.77 |
| State10 | 0.26 | 0.20 | 0.36 | 0.16 | 0.10 | 0.41 | 0.13 | 0.49 |
| State11 | 0.70 | 0.61 | 0.61 | 0.40 | 0.14 | 0.27 | 0.35 | 0.44 |
| State12 | 0.70 | 0.54 | 0.72 | 1.15 | 0.79 | 0.68 | 0.89 | 0.64 |
| State13 | 0.04 | 0.03 | 0.05 | 0.02 | 0.02 | 0.09 | 0.02 | 0.07 |
| State14 | 66.00 | 66.24 | 70.09 | 71.32 | 67.16 | 61.00 | 64.03 | 68.10 |
| State15 | 6.42 | 5.40 | 4.31 | 3.88 | 5.18 | 8.02 | 8.18 | 4.48 |
| State16 | 4.22 | 3.49 | 4.91 | 3.09 | 8.58 | 5.60 | 4.82 | 4.20 |
| State17 | 1.01 | 0.33 | 0.76 | 0.24 | 0.40 | 0.81 | 0.73 | 0.76 |
| State18 | 0.50 | 0.81 | 0.27 | 0.24 | 0.44 | 0.49 | 0.28 | 0.60 |

**Supplementary table1D:** abundance of different ChromHMM in neuroblastoma cell lines. Genic and intergenic regions, genome-wide.

|  | Short telomeres<br>Telomerase + |  |  |  | Long telomeres<br>Telomerase - |  |  |  |
| --- | --- | --- | --- | --- | --- | --- | --- | --- |
|  | IMR32 | NB5 | SKNBE2 | NBLS | CHLA90 | SKNMM | SKNFI | LAN6 |
| State1 | 0.56 | 0.27 | 0.51 | 0.63 | 0.62 | 0.64 | 0.52 | 0.61 |
| State2 | 0.01 | 0.00 | 0.02 | 0.01 | 0.01 | 0.03 | 0.01 | 0.03 |
| State3 | 0.51 | 0.58 | 0.74 | 0.40 | 0.57 | 0.51 | 0.54 | 0.78 |
| State4 | 0.96 | 1.28 | 1.78 | 1.28 | 0.66 | 1.21 | 1.13 | 1.42 |
| State5 | 1.00 | 1.79 | 0.83 | 1.34 | 1.32 | 1.33 | 0.87 | 0.91 |
| State6 | 1.93 | 3.65 | 2.70 | 2.73 | 1.83 | 2.25 | 1.74 | 2.78 |
| State7 | 0.15 | 0.26 | 0.30 | 0.10 | 0.21 | 0.35 | 0.21 | 0.17 |
| State8 | 6.97 | 7.26 | 5.88 | 7.41 | 7.37 | 8.08 | 7.14 | 6.32 |
| State9 | 3.19 | 2.11 | 1.46 | 1.73 | 0.69 | 2.46 | 3.53 | 2.68 |
| State10 | 0.18 | 0.14 | 0.27 | 0.12 | 0.07 | 0.29 | 0.10 | 0.35 |
| State11 | 0.57 | 0.46 | 0.50 | 0.33 | 0.11 | 0.21 | 0.27 | 0.34 |
| State12 | 0.54 | 0.40 | 0.56 | 0.90 | 0.60 | 0.51 | 0.67 | 0.49 |
| State13 | 0.03 | 0.02 | 0.04 | 0.02 | 0.01 | 0.06 | 0.01 | 0.05 |
| State14 | 70.52 | 70.69 | 73.04 | 74.79 | 70.69 | 65.90 | 68.02 | 71.91 |
| State15 | 6.28 | 5.64 | 4.30 | 3.85 | 5.04 | 7.77 | 8.13 | 4.30 |
| State16 | 5.10 | 4.26 | 6.05 | 3.87 | 9.34 | 7.08 | 6.09 | 5.54 |
| State17 | 0.99 | 0.38 | 0.73 | 0.24 | 0.43 | 0.83 | 0.73 | 0.71 |
| State18 | 0.51 | 0.82 | 0.28 | 0.26 | 0.44 | 0.49 | 0.28 | 0.61 |

**Table S2.**
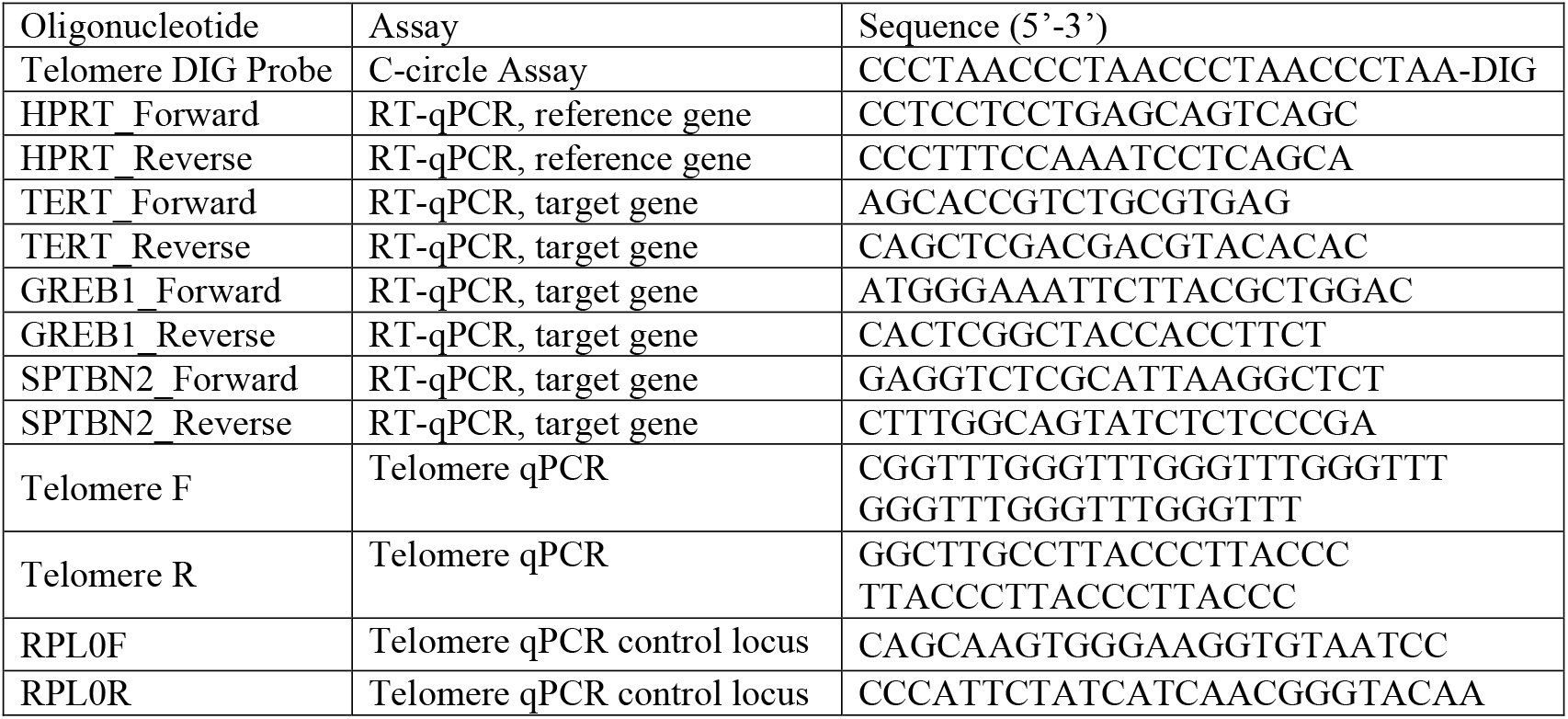
Sequences of oligonucleotides.

## References

1 Shay, J. W. & Wright, W. E. Telomeres and telomerase: three decades of progress. Nat Rev Genet 20, 299–309, doi:10.1038/s41576-019-0099-1 (2019).

2 Hemann, M. T., Strong, M. A., Hao, L. Y. & Greider, C. W. The shortest telomere, not average telomere length, is critical for cell viability and chromosome stability. Cell 107, 67–77, doi:10.1016/s0092-8674(01)00504-9 (2001).

3 Maciejowski, J. & de Lange, T. Telomeres in cancer: tumour suppression and genome instability. Nat Rev Mol Cell Biol 18, 175–186, doi:10.1038/nrm.2016.171 (2017).

4 Hanahan, D. & Weinberg, R. A. Hallmarks of cancer: the next generation. Cell 144, 646–674, doi:10.1016/j.cell.2011.02.013 (2011).

5 Zhang, J. M. & Zou, L. Alternative lengthening of telomeres: from molecular mechanisms to therapeutic outlooks. Cell Biosci 10, 30, doi:10.1186/s13578-020-00391-6 (2020).

6 Dagg, R. A. et al. Extensive Proliferation of Human Cancer Cells with Ever-Shorter Telomeres. Cell Rep 19, 2544–2556, doi:10.1016/j.celrep.2017.05.087 (2017).

7 Barthel, F. P. et al. Systematic analysis of telomere length and somatic alterations in 31 cancer types. Nat Genet 49, 349–357, doi:10.1038/ng.3781 (2017).

8 Mallepalli, S., Gupta, M. K. & Vadde, R. Neuroblastoma: An Updated Review on Biology and Treatment. Curr Drug Metab 20, 1014–1022, doi:10.2174/1389200221666191226102231 (2019).

9 Brodeur, G. M. Spontaneous regression of neuroblastoma. Cell Tissue Res 372, 277–286, doi:10.1007/s00441-017-2761-2 (2018).

10 Koneru, B. et al. Telomere Maintenance Mechanisms Define Clinical Outcome in High-Risk Neuroblastoma. Cancer Res 80, 2663–2675, doi:10.1158/0008-5472.CAN-19-3068 (2020).

11 Peifer, M. et al. Telomerase activation by genomic rearrangements in high-risk neuroblastoma. Nature 526, 700–704, doi:10.1038/nature14980 (2015).

12 Brodeur, G. M., Seeger, R. C., Schwab, M., Varmus, H. E. & Bishop, J. M. Amplification of N-myc in untreated human neuroblastomas correlates with advanced disease stage. Science 224, 1121–1124, doi:10.1126/science.6719137 (1984).

13 Schwab, M. et al. Amplified DNA with limited homology to myc cellular oncogene is shared by human neuroblastoma cell lines and a neuroblastoma tumour. Nature 305, 245–248, doi:10.1038/305245a0 (1983).

14 Schwab, M. et al. Enhanced expression of the human gene N-myc consequent to amplification of DNA may contribute to malignant progression of neuroblastoma. Proc Natl Acad Sci U S A 81, 4940–4944, doi:10.1073/pnas.81.15.4940 (1984).

15 He, S., Liu, Z., Oh, D. Y. & Thiele, C. J. MYCN and the epigenome. Front Oncol 3, 1, doi:10.3389/fonc.2013.00001 (2013).

16 Zeineldin, M. et al. MYCN amplification and ATRX mutations are incompatible in neuroblastoma. Nat Commun 11, 913, doi:10.1038/s41467-020-14682-6 (2020).

17 Yuan, X., Larsson, C. & Xu, D. Mechanisms underlying the activation of TERT transcription and telomerase activity in human cancer: old actors and new players. Oncogene 38, 6172–6183, doi:10.1038/s41388-019-0872-9 (2019).

18 Cheung, N. K. et al. Association of age at diagnosis and genetic mutations in patients with neuroblastoma. JAMA 307, 1062–1071, doi:10.1001/jama.2012.228 (2012).

19 Eustermann, S. et al. Combinatorial readout of histone H3 modifications specifies localization of ATRX to heterochromatin. Nat Struct Mol Biol 18, 777–782, doi:10.1038/nsmb.2070 (2011).

20 Lewis, P. W., Elsaesser, S. J., Noh, K. M., Stadler, S. C. & Allis, C. D. Daxx is an H3.3-specific histone chaperone and cooperates with ATRX in replication-independent chromatin assembly at telomeres. Proc Natl Acad Sci U S A 107, 14075–14080, doi:10.1073/pnas.1008850107 (2010).

21 Wong, L. H. et al. ATRX interacts with H3.3 in maintaining telomere structural integrity in pluripotent embryonic stem cells. Genome Res 20, 351–360, doi:10.1101/gr.101477.109 (2010).

22 Allis, C. D. & Jenuwein, T. The molecular hallmarks of epigenetic control. Nat Rev Genet 17, 487–500, doi:10.1038/nrg.2016.59 (2016).

23 Lewis, K. A. & Tollefsbol, T. O. Regulation of the Telomerase Reverse Transcriptase Subunit through Epigenetic Mechanisms. Front Genet 7, 83, doi:10.3389/fgene.2016.00083 (2016).

24 Dessain, S. K., Yu, H., Reddel, R. R., Beijersbergen, R. L. & Weinberg, R. A. Methylation of the human telomerase gene CpG island. Cancer Res 60, 537–541 (2000).

25 Takasawa, K. et al. DNA hypermethylation enhanced telomerase reverse transcriptase expression in human-induced pluripotent stem cells. Hum Cell 31, 78–86, doi:10.1007/s13577-017-0190-x (2018).

26 Stern, J. L. et al. Allele-Specific DNA Methylation and Its Interplay with Repressive Histone Marks at Promoter-Mutant TERT Genes. Cell Rep 21, 3700–3707, doi:10.1016/j.celrep.2017.12.001 (2017).

27 Esopi, D. et al. Pervasive promoter hypermethylation of silenced TERT alleles in human cancers. Cell Oncol (Dordr) 43, 847–861, doi:10.1007/s13402-020-00531-7 (2020).

28 Salgado, C. et al. Interplay between TERT promoter mutations and methylation culminates in chromatin accessibility and TERT expression. PLoS One 15, e0231418, doi:10.1371/journal.pone.0231418 (2020).

29 van Mierlo, G., Veenstra, G. J. C., Vermeulen, M. & Marks, H. The Complexity of PRC2 Subcomplexes. Trends Cell Biol 29, 660–671, doi:10.1016/j.tcb.2019.05.004 (2019).

30 Lee, D. D. et al. DNA hypermethylation within TERT promoter upregulates TERT expression in cancer. J Clin Invest 129, 223–229, doi:10.1172/JCI121303 (2019).

31 Nabetani, A. & Ishikawa, F. Unusual telomeric DNAs in human telomerase-negative immortalized cells. Mol Cell Biol 29, 703–713, doi:10.1128/MCB.00603-08 (2009).

32 Henson, J. D. et al. DNA C-circles are specific and quantifiable markers of alternative-lengthening-of-telomeres activity. Nat Biotechnol 27, 1181–1185, doi:10.1038/nbt.1587 (2009).

33 Qadeer, Z. A. et al. ATRX In-Frame Fusion Neuroblastoma Is Sensitive to EZH2 Inhibition via Modulation of Neuronal Gene Signatures. Cancer Cell 36, 512–527 e519, doi:10.1016/j.ccell.2019.09.002 (2019).

34 Farooqi, A. S. et al. Alternative lengthening of telomeres in neuroblastoma cell lines is associated with a lack of MYCN genomic amplification and with p53 pathway aberrations. J Neurooncol 119, 17–26, doi:10.1007/s11060-014-1456-8 (2014).

35 Dyer, M. A., Qadeer, Z. A., Valle-Garcia, D. & Bernstein, E. ATRX and DAXX: Mechanisms and Mutations. Cold Spring Harb Perspect Med 7, doi:10.1101/cshperspect.a026567 (2017).

36 Heaphy, C. M. et al. Altered telomeres in tumors with ATRX and DAXX mutations. Science 333, 425, doi:10.1126/science.1207313 (2011).

37 Blanco, E., Gonzalez-Ramirez, M., Alcaine-Colet, A., Aranda, S. & Di Croce, L. The Bivalent Genome: Characterization, Structure, and Regulation. Trends Genet 36, 118–131, doi:10.1016/j.tig.2019.11.004 (2020).

38 Kim, W. et al. Regulation of the Human Telomerase Gene TERT by Telomere Position Effect-Over Long Distances (TPE-OLD): Implications for Aging and Cancer. PLoS Biol 14, e2000016, doi:10.1371/journal.pbio.2000016 (2016).

39 Doheny, J. G., Mottus, R. & Grigliatti, T. A. Telomeric position effect--a third silencing mechanism in eukaryotes. PLoS One 3, e3864, doi:10.1371/journal.pone.0003864 (2008).

40 Robin, J. D. et al. Telomere position effect: regulation of gene expression with progressive telomere shortening over long distances. Genes Dev 28, 2464–2476, doi:10.1101/gad.251041.114 (2014).

41 Castelo-Branco, P. et al. Methylation of the TERT promoter and risk stratification of childhood brain tumours: an integrative genomic and molecular study. Lancet Oncol 14, 534–542, doi:10.1016/S1470-2045(13)70110-4 (2013).

42 Lee, D. D., Komosa, M., Nunes, N. M. & Tabori, U. DNA methylation of the TERT promoter and its impact on human cancer. Curr Opin Genet Dev 60, 17–24, doi:10.1016/j.gde.2020.02.003 (2020).

43 Renaud, S. et al. Dual role of DNA methylation inside and outside of CTCF-binding regions in the transcriptional regulation of the telomerase hTERT gene. Nucleic Acids Res 35, 1245–1256, doi:10.1093/nar/gkl1125 (2007).

44 Huang, F. W. et al. TERT promoter mutations and monoallelic activation of TERT in cancer. Oncogenesis 4, e176, doi:10.1038/oncsis.2015.39 (2015).

45 Zinn, R. L., Pruitt, K., Eguchi, S., Baylin, S. B. & Herman, J. G. hTERT is expressed in cancer cell lines despite promoter DNA methylation by preservation of unmethylated DNA and active chromatin around the transcription start site. Cancer Res 67, 194–201, doi:10.1158/0008-5472.CAN-06-3396 (2007).

46 Cohn, S. L. et al. Prolonged N-myc protein half-life in a neuroblastoma cell line lacking N-myc amplification. Oncogene 5, 1821–1827 (1990).

47 Li, Y. et al. Genome-wide analyses reveal a role of Polycomb in promoting hypomethylation of DNA methylation valleys. Genome Biol 19, 18, doi:10.1186/s13059-018-1390-8 (2018).

48 Cheng et al. Regulation of human and mouse telomerase genes by genomic contexts and transcription factors during embryonic stem cell differentiation. Sci Rep 7, 16444, doi:10.1038/s41598-017-16764-w (2017).

49 Brosnan-Cashman, J. A. et al. ATRX loss induces multiple hallmarks of the alternative lengthening of telomeres (ALT) phenotype in human glioma cell lines in a cell line-specific manner. PLoS One 13, e0204159, doi:10.1371/journal.pone.0204159 (2018).

